# Development and Comparative Evaluation of Endolysosomal Proximity Labeling-based Proteomic Methods in Human iPSC-derived Neurons

**DOI:** 10.1101/2020.09.15.298091

**Authors:** Ashley M. Frankenfield, Michael S. Fernandopulle, Saadia Hasan, Michael E. Ward, Ling Hao

**Affiliations:** Department of Chemistry, The George Washington University, Science and Engineering Hall 4000, 800 22^nd^ St., NW, Washington, DC, 20052, USA; National Institute of Neurological Disorders and Stroke, NIH, Bldg 35-2A, 35 Convent Dr. Bethesda, MD, 20892, USA

**Keywords:** Proximity labeling, Proteomics, Lysosome, Endolysosome, LAMP1, APEX, iPSC-derived neuron, Biotinylated proteins, Streptavidin

## Abstract

Proximity-based *in situ* labeling techniques offer a unique way to capture both stable and transient protein-protein and protein-organelle interactions. Combining this technology with mass spectrometry (MS)-based proteomics allows us to obtain snapshots of molecular microenvironments with nanometer resolution, facilitating the discovery of complex and dynamic protein networks. However, a number of technical challenges still exist, such as interferences from endogenously biotinylated proteins and other highly abundant bystanders, how to select the proper controls to minimize false discoveries, and experimental variations among biological/technical replicates. Here, we developed a new method to capture the proteomic microenvironment of the neuronal endolysosomal network, by knocking in (KI) an engineered ascorbate peroxidase (APEX) gene to the endogenous locus of lysosome-associated membrane protein 1 (LAMP1). We found that normalizing proximity labeling proteomics data to the endogenously biotinylated protein (PCCA) can greatly reduce variations and enable fair comparisons among different batch of APEX labeling and different APEX probes. We conducted comparative evaluation between this KI-LAMP1-APEX method and our two overexpression LAMP1-APEX probes, achieving complementary coverage of both known and new lysosomal membrane and lysosomal-interacting proteins in human iPSC-derived neurons. To summarize, this study demonstrated new analytical tools to characterize lysosomal functions and microenvironment in human neurons and filled critical gaps in the field for designing and optimizing proximity labeling proteomic experiments.

## INTRODUCTION

Protein-protein interactions are essential for almost all cellular processes and provide key insights into disease processes.^1,2^ Traditionally, protein-protein interactions are characterized by yeast two-hybrid assays or affinity-purification.^3–5^ However, affinity-purification often fails to capture transient and weak protein interactions, which are crucial for signal transduction and cellular transport processes. In recent years, proximity labeling (PL) techniques have emerged as a new class of tools for identifying both stable and transient protein interactions and molecular microenvironment in living cells and organisms.^6,7^ PL enzymes can be genetically fused to a target protein (i.e. “bait”) to biotinylate neighboring protein interactors (i.e. “prey”) upon activation. Biotinylated proteins can then be selectively purified and subsequently analyzed by mass spectrometry (MS)-based proteomics.

Biotin ligase and engineered ascorbate peroxidase represent the two most commonly used classes of PL enzymes, both developed in 2012.^8,9^ Bifunctional ligase/repressor (BioID) is a promiscuous mutant of *Escherichia coli* biotin protein ligase, which can be genetically tagged onto a bait protein to biotinylate neighboring proteins upon the addition of biotin substrate.^8^ However, BioID method often requires 12-24 hours of labeling, causing the diffusion of the reactive biotin cloud and non-specific labeling.^10^ Recently developed TurboID method mitigated this issue and shortened the labeling time to 10 min.^11^ The other type of PL enzyme is the engineered soybean ascorbate peroxidase (APEX), developed by the Ting group.^9,12^ Nowadays, APEX-based PL-MS has become a popular tool to capture protein-protein and protein-RNA interactions in different cellular compartments and whole organisms.^12–16^ The major advantages of APEX over biotin ligases are the rapid and highly efficient labeling reaction (≤ 1 min) and smaller size (27 kDa) compared to BioID/TurboID (35 kDa), minimizing the potential impact on the function of the fusion protein. Upon addition of the biotin-phenol substrate, the APEX enzyme can be activated by rapid H_2_O_2_ treatment to generate highly reactive phenoxyl radicals, linking biotin-phenol to the nearby proteins within a 10-20 nm radius. Biotinylated proteins can then be enriched by streptavidin-coated beads followed by MS-based proteomic strategies.

Despite the advances of PL methods, a number of technical challenges still exist. For example, the addition of biotin substrate in PL experiments enriches the endogenously biotinylated proteins within the mitochondria, which often represent the highest abundant biotinylated proteins in the dataset.^17^ The use of H_2_O_2_ for APEX activation raised concerns about potential oxidative stress.^18^ Experimental variations between different batches of cell culture and PL labeling undermine the accurate and reproducible quantification of protein-protein interactions. The selection of proper controls is also critical to minimize both false-positive and false-negative protein interactions. These challenges have limited the accuracy of PL-MS and generated technical gaps between method development and successful biological applications.

In this study, we aimed to address these major challenges by developing a new endogenous APEX-based PL-MS method and conducting a thorough investigations of key factors involved in the PL-based proteomic workflow. We knocked in (KI) the APEX gene onto the endogenous locus of lysosomal-associated membrane protein 1 (LAMP1), localizing to the cytosolic surface of endolysosomes in human induced pluripotent stem cells (hiPSCs)-derived neurons, namely KI-LAMP1-APEX. Lysosomes are acidic vacuoles that digest and recycle macromolecules through a cellular mechanism known as autophagy.^19^ Lysosomes are particularly important for neurons, because neurons are highly polarized, postmitotic, and rely on autophagy to maintain cellular homeostasis.^20^ LAMP1 is an abundant transmembrane protein on the endolysosomal membrane and has served as a classical lysosomal marker.^21^ The LAMP1-APEX proteomic workflow was systematically optimized to reduce interferences and variations, improve reproducibility and specificity, and address the potential effect of H_2_O_2_ treatment. A comparative evaluation was also conducted between KI-LAMP1-APEX method and our two other overexpressed LAMP1-APEX probes to discover known and novel lysosomal membrane and lysosomal-interacting proteins in hiPSC-derived neurons.

## EXPERIMENTAL SECTION

### Human iPSC Culture and Development of LAMP1-APEX Lines

Wild type hiPSCs from a WTC11 control male were obtained from Coriell and routinely maintained in Matrigel-coated plates in Essential 8 Flex media as described previously.^22^ Three stable LAMP1-APEX iPSC lines were generated in this study for comparative analysis, endogenous knock-in (KI) LAMP1-APEX, KuB-LAMP1-APEX (high overexpression) and KuD-LAMP1-APEX (modest overexpression). The expression levels of LAMP1-APEX in iPSCs are KuB> KuD> KI. Nuclear exporting signal (NES)-APEX (overexpression) was also generated as a cytosolic localized APEX line. The detailed steps to generate and validate stable APEX iPSCs lines are described in Supporting Information.

### Differentiation of hiPSCs into i^3^Neurons

We implemented the advanced i^3^Neuron platform developed in the Ward Lab as our primary cellular platform in this study.^22,23^ The hiPSCs used in this study contain a stably integrated doxycycline-inducible neurogenin-2 cassette, which promotes the differentiation of the stem cells into functionally mature glutamatergic cortical neurons in two weeks.^23^ LAMP1-APEX hiPSCs were differentiated into cortical neurons as described previously.^22^ Briefly, transgenic hiPSCs were seeded into a 15cm dish with Neuronal Induction Medium^22^, which promotes terminal differentiation within three days. Then, the cells were dissociated with Accutase (StemPro), seeded onto polyornithine-coated plates, and maintained in Cortical Neuron Culture Medium^22^. Half-medium change was conducted every other day until neuronal maturation for APEX activation experiment and harvest.

### In vivo Proximity Labeling in hiPSC-derived Neurons

Prior to APEX activation, neurons were incubated with 500 μM biotin-phenol at 37 °C for 30 min. Then, 1 mM of H_2_O_2_ was added to the cells for exactly 1 min at 37 °C to activate APEX enzyme. The PL reaction was immediately terminated by aspirating the growth medium and rapid washing and incubating with ice-cold quench buffer (10mM sodium azide, 10mM sodium ascorbate, 5mM TROLOX in PBS) before lysing the neurons in ice-cold lysis buffer (50 mM Tris-HCl, 500 mM NaCl, 0.2 % SDS, 1mM DTT, 10mM sodium azide, 10 mM sodium ascorbate, 5 mM TROLOX, cOmplete mini protease inhibitor). Neurons were then rocked at 4 °C for 10 min, scraped into QSonica (Q800R) sonication tubes, and sonicated for 15 min at 2 °C with alternating 40 s on, 20 s off cycles. The lysate was centrifuged at 4 °C at 16,500 g to collect supernatants and stored at −80 °C.

### Beads Titration Assay to Determine the Optimal Streptavidin Beads/Proteins Ratio

Biotinylated proteins can be captured by streptavidin-coated beads because of the strong non-covalent interactions between biotin and streptavidin (Kd ~10^-14^). The optimal beads/protein ratio can be determined by the bead titration assay. Protein concentrations of neuron lysates were first determined by DC (detergent-compatible) Colorimetric Protein Assay (Bio-Rad). Same amount of protein lysate were added to each 0.5 mL tube containing a series volume of streptavidin magnetic beads (GE) (0, 0.5, 1, 2, 5, 10, 20, 40 μL), followed by overnight rotation at 4 °C. The next morning, the tubes were placed on a magnetic rack, and 2 μl of the supernatant from each tube was spotted on a dry nitrocellulose membrane. Once completely dried, the membrane was incubated in Odyssey Blocking Buffer for 1 hour, and then Streptavidin Alexa Fluor 680 conjugate (1:1000 in blocking buffer) for 1hr and washed 5 times with TBST buffer. The fluorescent signal of each dot on the membrane was measured under 700 nm wavelength.

### Biotinylated Protein Pull Down and On-Bead Digestion

Protein lysate samples were incubated with streptavidin magnetic beads (with optimal protein/beads ratio) overnight at 4 °C. Supernatants were then removed and beads were washed twice with each of the four sequential buffers (Buffer A: 2% SDS; Buffer B: 50mM Tris-HCl, 500mM NaCl, 0.1% deoxycolic acid, 1% Triton-X, 1 mM EDTA; Buffer C: 50mM Tris-HCl, 250mM NaCl, 0.5% deoxycolic acid, 0.5% NP-40, 1 mM EDTA; Buffer D: 50mM Tris-HCl, 2 M Urea). Magnetic beads were washed two more times with Buffer D to remove all residue detergents. To reduce the disulfide bonds, 5 mM of Tris(2-carboxyethyl)phosphine (TCEP) was added to the beads (resuspended in 100 μL of Buffer D) for 30 min, 15 mM of iodoacetamide for 30 min, and 5 mM of TCEP for 10 min in a ThermoMixer shaking at 1200 rpm at 37 °C. Trypsin/Lys-C Mix enzyme (Promega) was then added to the beads for a 16-hour on-beads digestion at 37 °C in a ThermoMixer shaking at 1200 rpm. Then a half amount of Trypsin/LysC mix was added for an additional 3 hours of digestion. The sample tubes were then briefly centrifuged and put on a magnetic rack to collect supernatants. The magnetic beads were washed with 50 μL of 50 mM Tris buffer, and supernatants were combined. To quench the digestion, 10% trifluoroacetic acid was added to the supernatant until pH <3. Peptide desalting was achieved by Waters Oasis HLB 96-well extraction plate based on manufacture protocol. Peptide samples were dried under SpeedVac and stored at −80°C.

### LC-MS Analysis

Dried peptide samples were resuspended in 10 μL of 2% acetonitrile (ACN), 0.1% formic acid (FA) in LC-MS grade water and centrifuged to collect supernatant. LC-MS analyses were performed on a Dionex UltiMate 3000 RSLCnano system coupled with a Thermo Scientific Q-Exactive HF mass spectrometer. The mobile phase buffer A was 0.1% FA, 5% DSMO in H_2_O, and buffer B was 0.1% FA, 5% DSMO in ACN. Two microliters were injected for each sample onto an Easy-spray PepMap C18 column (2 μM, 100 Å, 75 μM ×75 cm) with a 2-hour LC gradient and 60 °C column temperature. The LC flow rate is 0.2 μl/min. MS was scanned from *m/z* 360 to 1500 at 120K resolution in top 15 data dependent acquisition. Parent masses were isolated (m/z 1.4 window) and fragmented with higher-energy collisional dissociation (HCD) with a normalized collision energy of 27% and dynamic exclusion time of 22.5s. Maximum injection times were 30 ms for MS and 35 ms for MS/MS. Automatic gain control (AGC) targets were 1×10^6^ for MS and 2×10^5^ for MS/MS. An exclusion list was used with interference peptides obtained from digesting streptavidin beads (no input protein) with an exclusion mass tolerance of 5 ppm and retention time window of 2 min.

### Proteomics Data Analysis

LC-MS/MS raw files were analyzed by MaxQuant (1.6.10.43) software for peptide/protein identification and quantification.^24^ Uniprot *Homo sapiens* proteome database (Swiss-Prot, cononical) was used for protein identification (1% false discovery rate cutoff) with a fixed modification of cysteine carbamidomethylation and variable modifications of oxidation of methionine, acetylation of protein N-terminus, and biotin-phenol modification of tyrosine. PEAKS Studio X software was used to conduct a semi-open post-translational modification (PTM) search. The PEAKS PTM algorithm was used to search for 309 potential pre-set modifications. Additional modifications for oxidation at other amino acid residues were imported from the Unimod database.^25^ Maxquant and PEAKS output files were analyzed in Excel and R for statistical analysis. Protein network analysis was conducted with STRING.^26^ For data normalization to the most abundant biotinylated protein (PCCA), raw protein intensities from Maxquant output were normalized to the PCCA intensities followed by log_2_-transformation before statistical analysis. Raw proteomics data from this manuscript are available through the MassIVE repository^27^ (Identifier: MSV000086260).

### Fluorescence Imaging of Neurons

Human i^3^Neurons expressing LAMP1-APEX were incubated in 500 μM phenol-biotin at 37 °C for 30 min, stimulated with 1mM H_2_O_2_ for one second, and immediately fixed in 4% paraformaldehyde. Fixed neurons were washed 3 times in PBS, blocked and permeabilized (3% donkey serum and 0.1% saponin in PBS) for 30 min at room temperature (RT), and incubated with primary anti-LAMP1 antibody (mouse monoclonal H3A4, DHSB, 1:1000) overnight on a nutator at 4 °C. The following day, fixed neuron dish were washed 3 times with PBS, incubated with anti-mouse AF561 secondary antibody (Jackson ImmunoResearch) and Streptavidin-680 (1:1000) for 1 hour at RT, washed twice with PBS, incubated with Hoechst (nuclear marker) for 10 min at RT, and final PBS wash twice. Fixed neurons were visualized with a Nikon Eclipse Ti spinning disk confocal microscope.

## RESULTS AND DISCUSSIONS

### Development and Validation of Endogenous KI-LAMP1-APEX Probe

Lysosomes frequently make contacts with other organelles for autophagy and other cellular processes, but these transient and dynamic interactions have historically been restricted to individual observations by live cell microscopy.^20^ In order to study both stable and transient lysosomal interactions in a high-throughput fashion, we developed a new endogenous LAMP1-APEX probe in human iPSC-neurons. The overall workflow of LAMP1-APEX proteomics is illustrated in Figure 1A. Human iPSCs expressing the LAMP1-APEX fusion protein were differentiated into cortical neurons in 2 weeks based on our i^3^Neuron protocol.^22^ To activate the APEX labeling, neurons were incubated with biotin-phenol for 30 min and H_2_O_2_ for 1 min. APEX enzyme catalyzed the reaction by generating highly reactive phenoxyl radicals that form covalent bonds between biotin-phenol and electron-rich amino acid residues (e.g. tyrosine) from proteins within a 10-20 nm labeling radius (Figure 1B).

**Figure 1.**
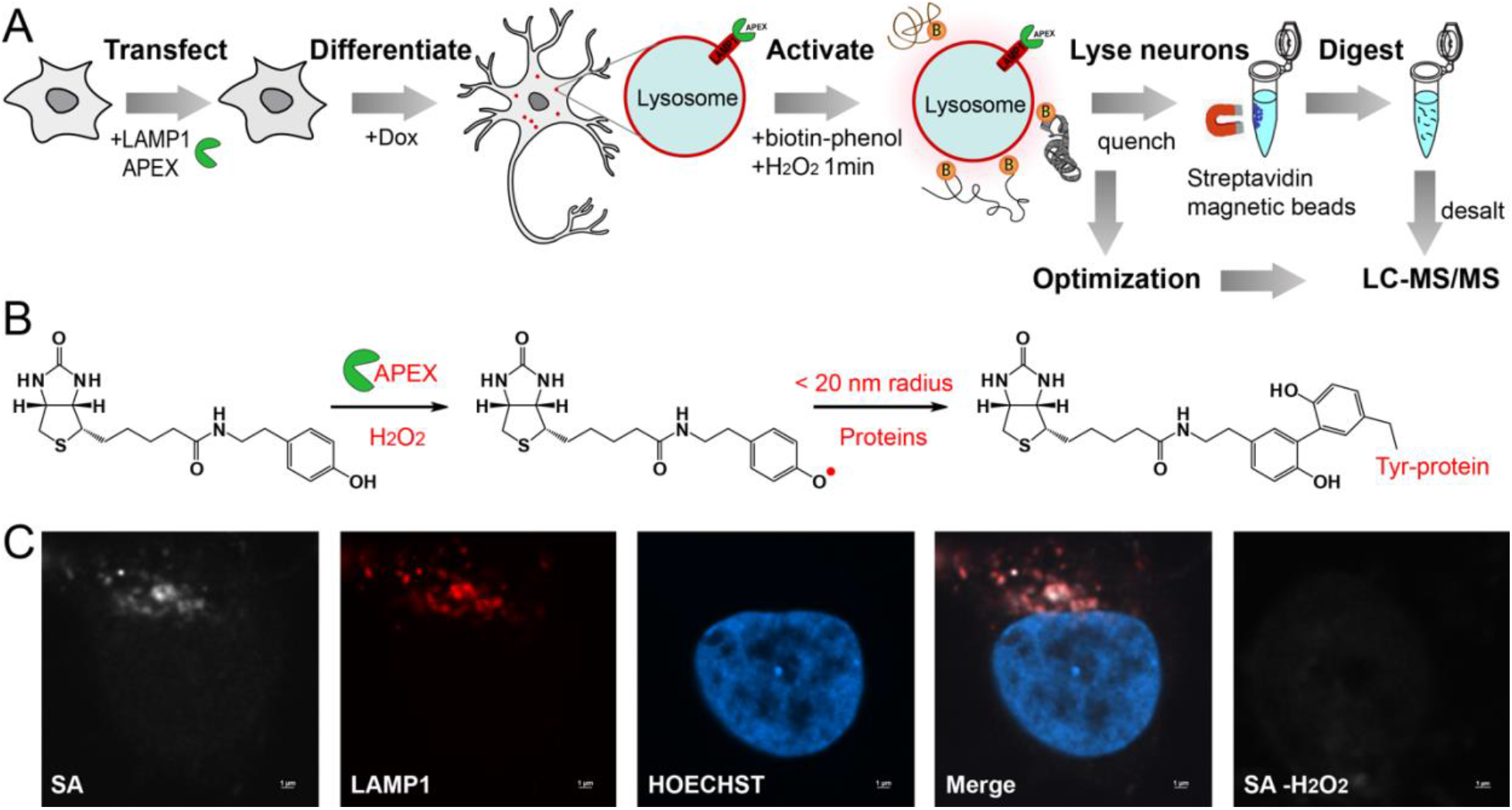
Development of endogenous lysosomal proximity labeling proteomics. (A) Illustration of LAMP1-APEX proteomic workflow in human iPSC-derived neurons. (B) Schematic of APEX enzymatic labeling reaction. (C) Fluorescence imaging of KI-LAMP1-APEX activity in an iPSC-derived neuron. Biotinylation is visualized by staining against streptavidin (SA) Fluor 680 (far red). LAMP1 is an endolysosome marker (red). Hoechst is a nuclear marker (blue).

In our recent study about lysosomal trafficking in neurons, we overexpressed KuB-APEX tagged LAMP1 protein in hiPSC-derived neurons and identified Annexin A11 as a molecular tether that links RNA granules to lysosomes during long-distance RNA transport in neuronal axons.^28^ Although overexpressing proteins is a common practice in cell biology, it might generate overexpression artifacts that can be picked out by the sensitive LC-MS platform. For instance, overexpressing LAMP1 might expand a small percentage of its localization to the plasma membrane, increasing non-specific labeling background.^29^

To address this issue, we developed a new LAMP1-APEX probe by knocking in (KI) APEX to the endogenous LAMP1 locus, namely KI-LAMP1-APEX. The correct localization of LAMP1-APEX probe was confirmed by fluorescence microscopy. As shown in Figure 1C **and Figure S1**, biotinylated protein signals stained by streptavidin (SA) exhibit excellent colocalization with LAMP1-staining in neurons. In the negative control without H_2_O_2_ activation, no biotinylation signal was observed. The APEX activity was also validated in a dot-blot assay (Figure 2A). The neuronal protein lysate (50 μg) was spotted onto a nitrocellulose membrane and stained against SA. The APEX enzyme was only activated when exposed to both biotin-phenol and H_2_O_2_. In control neurons with no APEX expression, no clear signals were observed. Therefore, our KI-LAMP1-APEX probe demonstrated specific endolysosomal localization and biotinylation in hiPSC-derived neurons.

**Figure 2.**
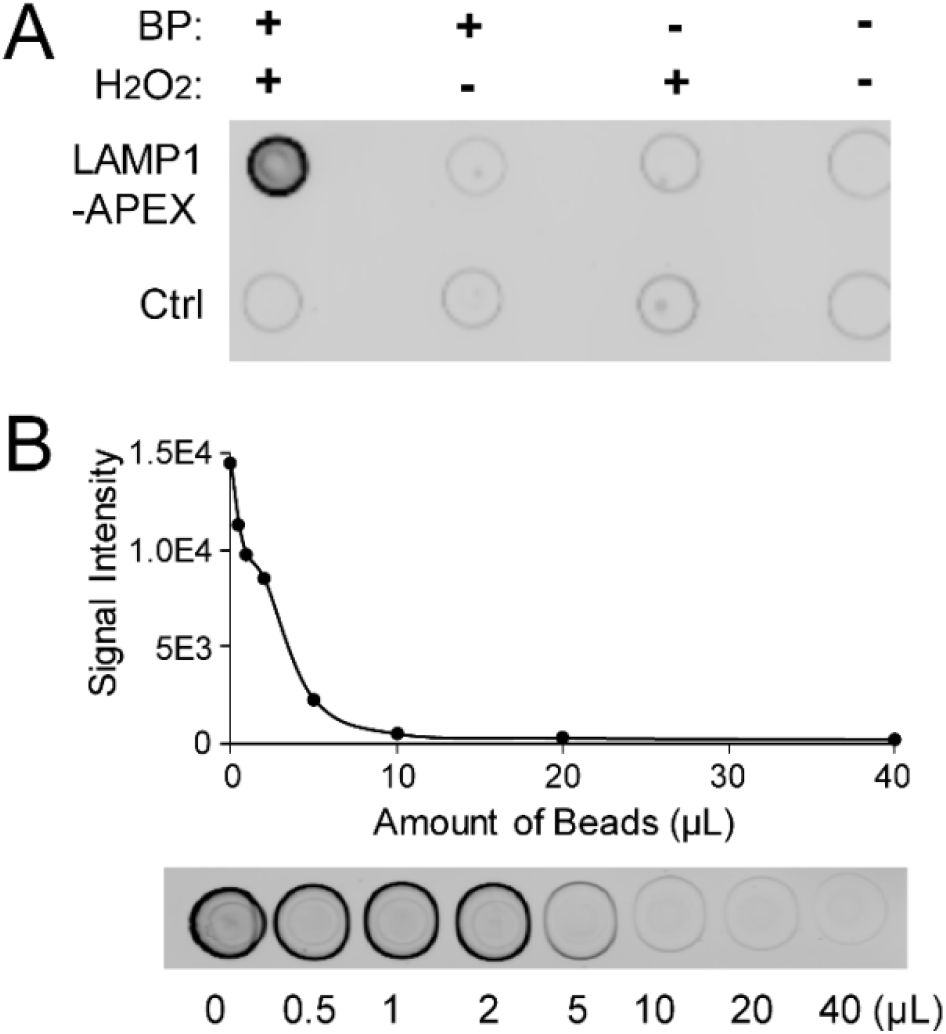
Validation and optimization of KI-LAMP1-APEX probe with streptavidin dot-blots. (A) Streptavidin dot-blot validation of APEX activity. (B) Beads titration assay to determine the optimal beads/proteins ratio.

### Optimization of the Proximity Labeling Experiment

Despite various applications of PL-proteomics in biological studies, technical challenges still exist that undermine the specificity and accuracy in identifying and quantifying protein interactions. Here, we conducted a systematic optimization of the key factors involved in PL-proteomics to achieve the optimal beads-to-protein ratio, improved reproducibility, reduced interferences, and optimal digestion efficiency.^30^

To ensure the complete and reproducible capture of all biotinylated proteins, streptavidin-coated beads are often added in excess. However, excess beads can cause overwhelming signals of streptavidin peptides from on-beads digestion. We conducted a beads titration assay to determine the optimal beads/proteins ratio. The dot-blot signals from supernatant of beads-protein mix are directly proportional to the amount of biotinylated proteins that are not captured by the beads. As illustrated in Figure 2B, fluorescence signals of the dot-blots declined with an increase amount of beads until reaching a plateau at ~5 μL of beads per 50 μg of input protein lysate, indicating the complete capture of biotinylated proteins from the sample. Therefore, the optimal beads/proteins ratio for KI-LAMP1-APEX proteomics was ~0.1 μL of beads per μg of input proteins. The optimal beads/proteins ratio is highly dependent on the bait protein of interest, and therefore beads titration assay is necessary for each PL probe targeting on different bait protein of interest or has different expression level of the bait protein.

To evaluate the interference background signals from on-beads digestion, we digested 200 μL of streptavidin magnetic beads without any input protein with 2 μg of Trypsin/LysC enzyme. As shown in Figure 3A, LC-MS chromatography was overwhelmed with streptavidin peptides from the beads. A total of 35 interference proteins and 384 peptides were identified originated from streptavidin, trypsin, keratins, *et.al*. When compared to an actual APEX proteomic experiment, these interference signals accounted for 2% of the identified proteins, 3% of peptides, and 8% of MS/MS scans. Based on these interference peptides, we generated an MS/MS exclusion list to be used for all PL experiments (**Table S1**). The starting *m/z* for full MS scan was also adjusted from 350 to 360 to exclude several abundant interference peaks (e.g. *m/z* 354.7068 streptavidin peptide). An alternative to on-beads digestion is to use anti-biotin antibodies to enrich biotinylated proteins, which can be eluted from anti-biotin antibody-coated beads and preserve the biotinylation sites on proteins, but with potential non-specific binding issue from the antibody.^31,32^ Excess biotin in harsh detergent buffer with heating can also be used to elute biotinylated proteins from streptavidin beads, but detergent must be removed before LC-MS analysis.^33^

**Figure 3.**
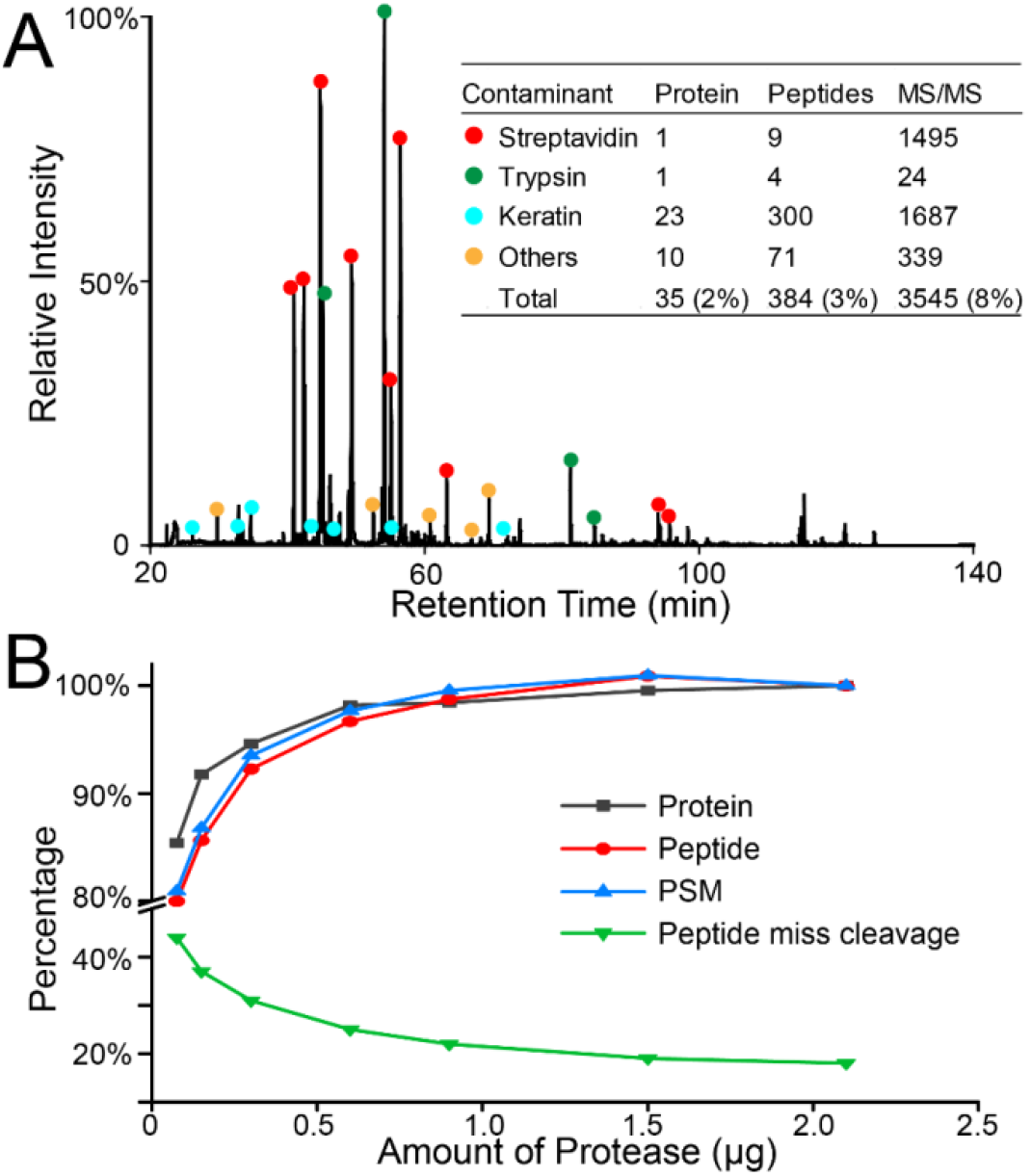
Evaluation of interference signals and different amount of proteases for on-beads digestion. (A) Base peak chromatogram of digesting streptavidin magnetic beads with no input proteins. The interference percentages were generated in comparison with an actual APEX proteomics experiment. (B) Optimization of the amount of Trypsin/LysC mix for on-beads digestion with 250 μL of streptavidin magnetic beads and corresponding input protein amount based on beads-titration assay.

Although total protein concentration was determined after cell lysis, the concentration of enriched proteins by streptavidin beads is unknown. To achieve the optimal on-beads digestion efficiency and minimum interferences from trypsin, we optimized the amount of protease for on-beads digestion. Compared to trypsin alone, Trypsin/Lys-C mix resulted in 15% more protein and peptide IDs with much less miscleavages (data not shown), in agreement with previous findings.^34^ As shown in Figure 3B, with an increased amount of Trypsin/Lys-C mix, the number of identifications increased and peptide miscleavages decreased until reaching a plateau at ~1-1.5 μg of protease per 250 μL of streptavidin magnetic beads (GE). Most identified peptides had +2 or +3 charges, and increased amount of protease shifted the peptides towards lower charges and smaller precursor masses (**Figure S2**).

### Controlling the false discoveries in proximity labeling proteomics

An inherent challenge in identifying protein interactions is the false discoveries, both false positives and false negatives.^35^ For PL-proteomics, false positive interactions include proteins that bind non-specifically to beads, endogenously biotinylated proteins, and other highly abundant bystanders. False negative interactions include low abundance and transient/weak protein interactions which can be ruled out if using overly strict cut-off thresholds. Here, we demonstrated that rational experimental design with data normalization and proper selection of controls can reduce these false positive and false negative discoveries.^30,36^

The presence of highly abundant endogenously biotinylated proteins has been a major concern in PL-proteomics.^18^ Within the mitochondria, pyruvate carboxylase (PC), 3-methylcrontonyl-coA carboxylase (MCC), propionyl-CoA carboxylase (PCC), and acetyl-CoA carboxylase 1 (ACACA) are known to be endogenously biotinylated.^37^ Upon the addition of biotin substrate, these endogenously biotinylated proteins as well as their interacting proteins in the mitochondria can be enriched by streptavidin beads, accounting for a major source of false positives in PL-MS. The heatmap intensities of endogenously biotinylated proteins in our APEX datasets are shown in Figure 4A. Although these abundant proteins cannot be removed from the samples, we found that normalizing the proteomic dataset to the most abundant endogenously biotinylated protein (PCCA) can greatly reduce variations for biological replicates in the same batch of APEX labeling as well as different batches of APEX experiments (1 month apart) (Figure 4B, **Figure S3**). Normalization to PCCA improved the reproducibility between replicates and allowed the fair comparison among different batches of APEX experiments and different APEX probes.

**Figure 4.**
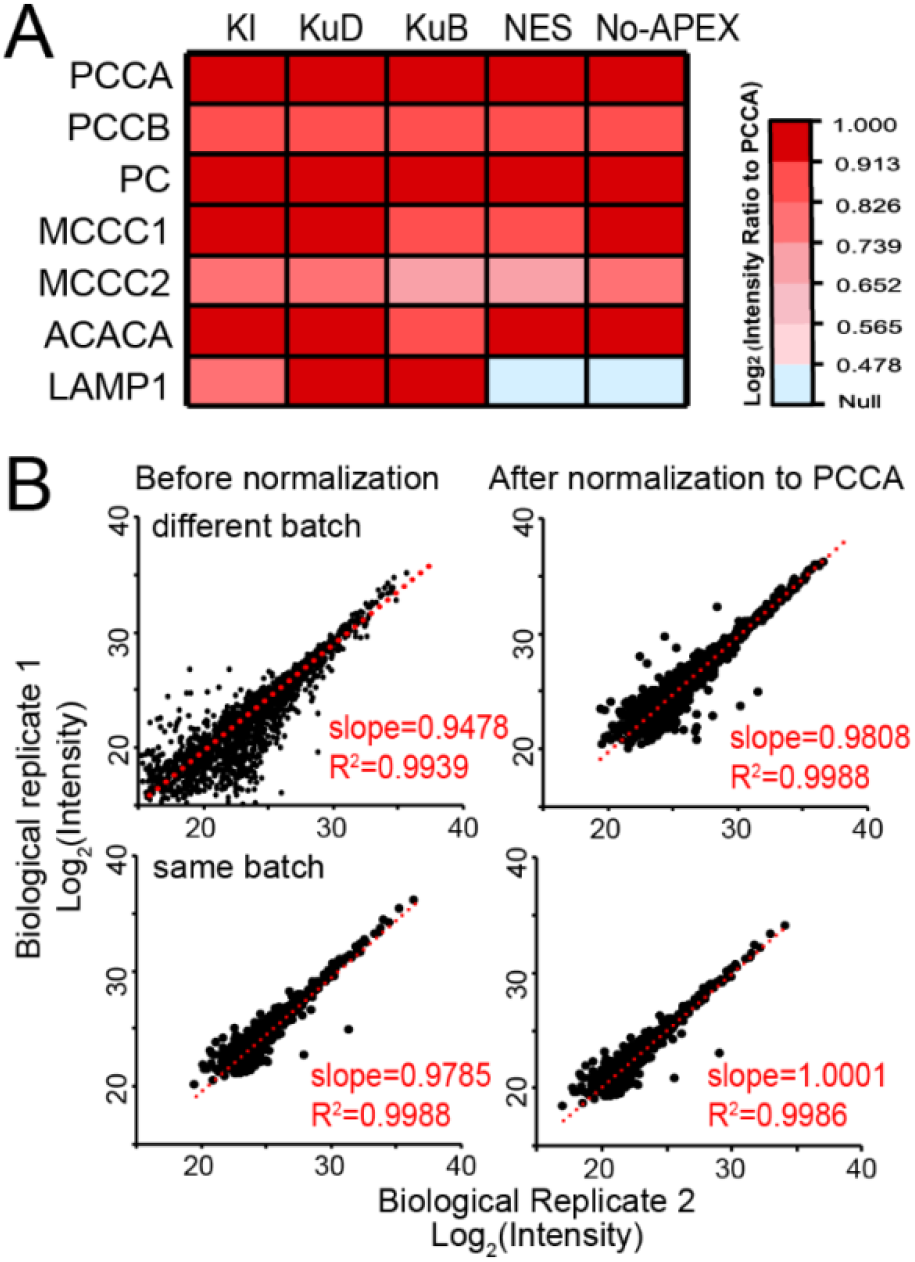
Normalization of APEX proteomics data to the endogenously biotinylated protein, PCCA. (A) Heatmap intensities of endogenously biotinylated proteins and LAMP1 bait protein across different APEX probes and controls. (B) Intensity scatter plots between biological replicates of KI-LAMP1-APEX data before and after normalization to PCCA for different batch of APEX labeling (upper panel) and same batch of APEX labeling (lower panel). Slope and R^2^ of 1 represent perfect reproducibility.

Designing/selecting the appropriate control group is critical in controlling false discoveries in PL-MS studies. We established two control lines for our LAMP1-APEX neurons: cytosol-localized NES-APEX line, and parental line that has no APEX expression. As illustrated in Figure 5 and **Figure S4**, all quantified proteins are ranked based on their abundance ratio between LAMP1-APEX vs. control lines. For KI neurons, the parental line lacking APEX is a better control (Figure 5A). Using NES-APEX control for KI neurons caused massive false negatives, which could be due to the disparity in APEX expression levels (Figure 5B). Whereas, for high expression KuD/KuB neurons, the line lacking APEX created substantial false positives (Figure 5C, **Figure S4A**). Thus, NES-APEX line is a better control for overexpression LAMP1-APEX probes (Figure 5D, **Figure S4B**).

**Figure 5.**
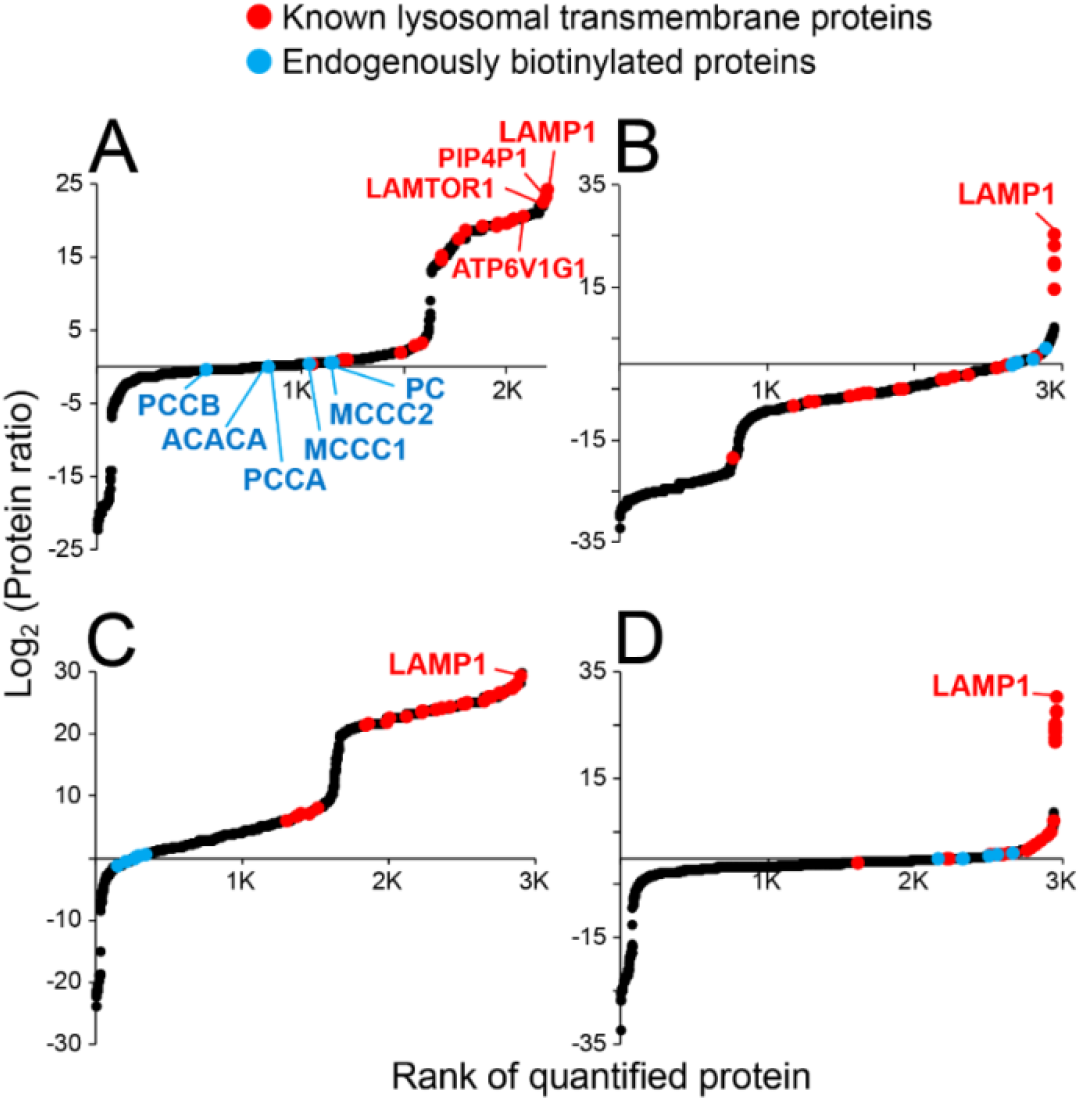
Evaluation of false discoveries in LAMP1-APEX proteomics using different control datasets. Scatter plots of the rank of protein ratios in (A) KI-LAMP1-APEX Proteomics with No-APEX parental line as control; (B) KI-LAMP1-APEX with NES-APEX as control; (C) KuD-LAMP1-APEX with No-APEX as control; (D) KuD-LAMP1-APEX with NES-APEX as control. Protein intensities were normalized to endogenously biotinylated protein, PCCA, and averaged among four biological replicates.

APEX activation requires brief H_2_O_2_ (1 mM for 1 min) and biotin-phenol (500 μM for 30 min) treatment for cells, which raised the concerns for possible oxidative stress and perturbation of cellular processes. To address this question directly, we conducted whole cell lysate proteomics from parental iPSC-neurons (no-APEX) under different treatments (H_2_O_2_ treated vs. biotin-phenol treated vs. control). Label-free proteomics quantification (N=3) identified 4339 proteins from neurons, but none of them reached statistical significance (Figure 6A, **Table S2**). Interestingly, glutathione peroxidase 4 (GPX4) and mitogen-activated protein kinase 3 (MAPK3), which function to protect against oxidative stress, were upregulated in H_2_O_2_ vs. control group without reaching statistical significance.^38,39^ To evaluate possible PTMs caused by H_2_O_2_ and biotin-phenol, we conducted a semi-open PTM search for the proteomics dataset with more than 300 PTMs from PEAKS software and Unimod database.^25^ As expected, methionine was found to be the most oxidized amino acid residue in peptides/proteins (Figure 6B). No obvious increase or uncommon PTMs was observed in H_2_O_2_ and biotin-phenol treatment groups vs. control, except that H_2_O_2_ treated group did present more numbers of oxidized peptide compared to controls. Therefore, the activation steps for APEX did not exert significant impact on cellular proteome other than enriching the endogenously biotinylated proteins and slightly higher number of oxidized peptides. However, a strict control of the concentration and time for H_2_O_2_ treatment is critical to minimize oxidative stress.

**Figure 6.**
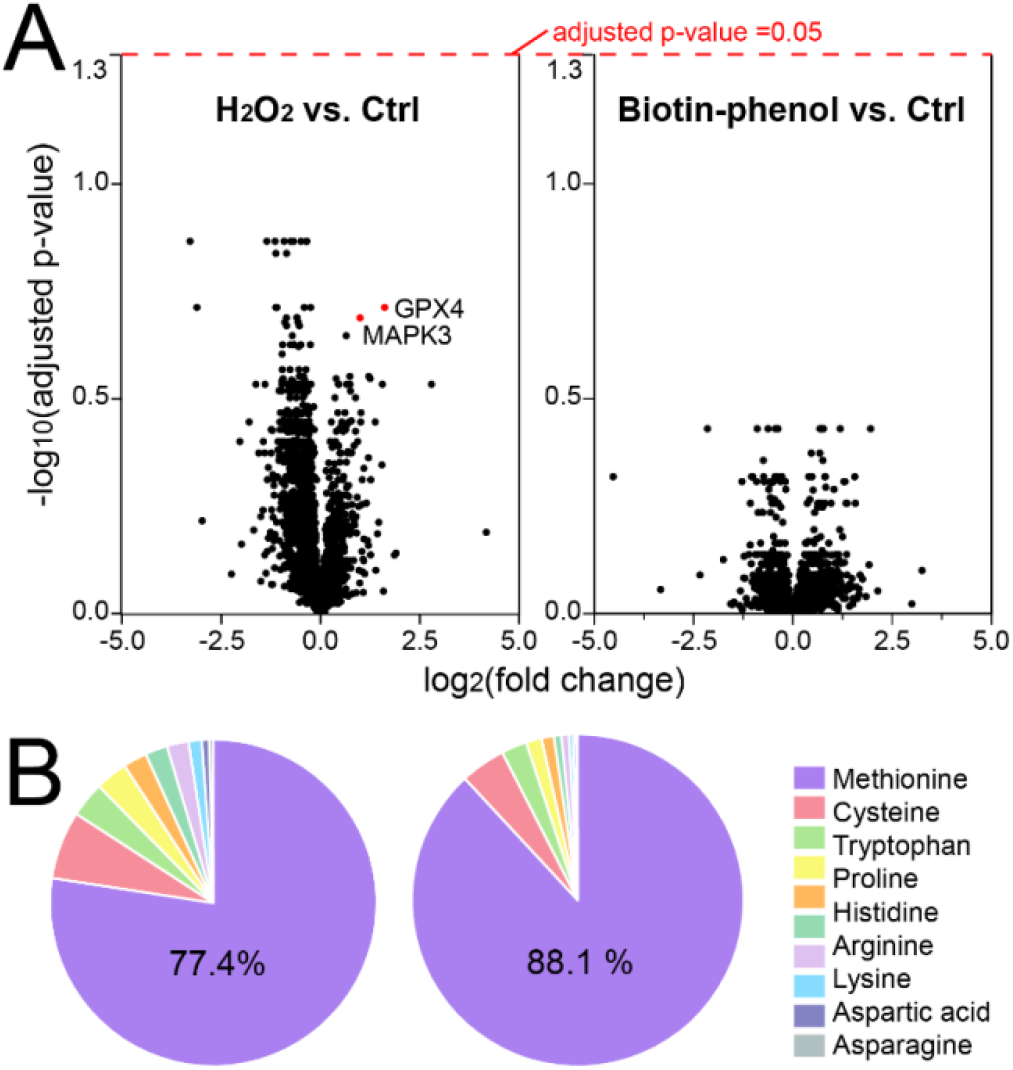
Evaluation of potential influence of H_2_O_2_ and PB treatment to the whole cell proteome. (A) Volcano plots of all the quantified proteins from H_2_O_2_ vs. Ctrl and PB vs. Ctrl groups in the whole cell protein extract. (B) Pie graph of oxidized amino acid residues from the identified proteins in iPSC-derived neurons.

### Comparison of Different Lysosome Proximity Labeling Probes

Different types of APEX probes have been developed for various biological applications, but their proteomic performances have rarely been compared. Here, we comparatively evaluated our three LAMP1-APEX probes, KI (endogenous expression), KuB (overexpression), and KuD (modest overexpression). The proteomic profiles are compared in Figure 7A. The KuB/KuD overexpression probes led to higher total numbers of identified proteins but also slightly higher false positives (e.g. nucleus and plasma membrane proteins) compared to the endogenous KI probe. The KuB neurons had slightly more IDs than KuD. These results were validated by the beads titration assay, where the optimal amount of beads per μg of input proteins was determined to be 0.1, 0.25, and 0.25 for KI, KuB, and KuD neurons, respectively (Figure 2B and **Figure S5**). Because the KI probe is expressed at physiological level, the proteomic results are less prone to false positives and mislocalization of LAMP1. However, at least 2.5 folds more starting material is needed for the KI neurons compared to KuB/KuD. Therefore, scalability of the cell lines and overexpression artifacts must be considered when determining the appropriate probe for PL experiments. The complete protein list and comparisons from all APEX probes are provided in **Table S3** and **Figure S6**.

**Figure 7.**
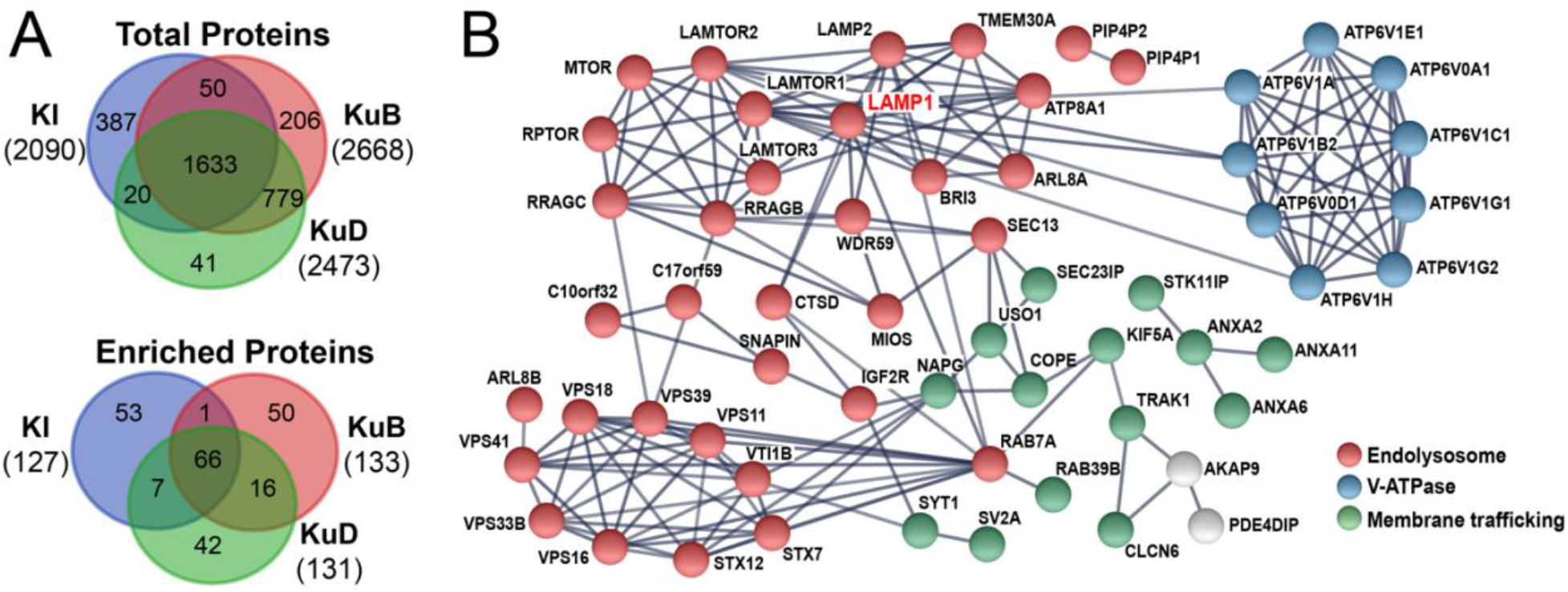
Comparative evaluation of different endolysosome proximity labeling probes. (A) Venn diagram of all identified proteins and truly enriched proteins after ratiometric analysis vs. controls from KI, KuB, and KuD LAMP1-APEX iPSC-derived neurons. (B) STRING protein network of overlapped proteins from three LAMP1-APEX probes.

After comparing to proper controls (KI vs. No-APEX control; KuB/KuD vs. NES-APEX control), all three of our LAMP1-APEX methods successfully identified and quantified specific endolysosomal membrane and membrane interacting proteins with complementary coverage (Figure 7A, **Table S4**). Protein network analysis was conducted for the overlapping proteins from at least two LAMP1-APEX methods (Figure 7B). These proteins are stable lysosomal membrane proteins (e.g. LAMP1, LAMP2, LAMTORs, PIP4P1, PIP4P2, V-ATPases) as well as transient lysosomal interacting proteins related to lysosomal transport, mobility and signaling pathways. For instance, the identified Rab GTPases (Rab7 and Rab39) and SNARE protein family (STX7, STX12, VTI1B) play key roles in autophagy, endolysosomal trafficking, and lysosome biogensis.^40,41^ The homotypic fusion and vacuole protein sorting (HOPS) complex is composed of six vacuolar protein sorting (Vps) subunits that act as a tether between the autophagosome and lysosome for autophagy and interact with the Rab GTPases.^40,41^ Multiple components of the V-ATPase complex were identified which maintains the acidic environment of the lysosomes and regulates the nutrient sensing and amino acid efflux through the mTORC1 pathway.^42^ Key proteins in the mTORC1 pathway (LAMTOR1,2,3, RRAGC, RRAGB, RPTOR, MTOR) were also identified in our LAMP1-APEX proteomic experiments.

## CONCLUSIONS

We developed a new endolysosomal proximity labeling proteomic method, KI-LAMP1-APEX, to study the endolysosomal membrane and membrane interacting proteins in human iPSC-derived neurons. Several key technical challenges in the proximity labeling field were addressed here. We recommend normalizing PL proteomics data to the endogenously biotinylated protein (PCCA), optimizing protein/beads ratio and protease amount, and selecting proper controls for each PL probe to reduce variation and false positive/negative interactions. Hydrogen peroxide treatment during APEX activation did not cause significant proteome perturbation but the time and concentration of H_2_O_2_ treatment need to be strictly controlled. Furthermore, comparative evaluations of different LAMP1-APEX probes (KI, KuD, and KuB) revealed complementary proteome coverage and provided key references for the rational design of proximity labeling experiments. To summarize, our study established new endolysosomal proximity labeling methods that can serve as promising tools to study lysosomal functions and filled critical gaps in the field of proximity labeling and affinity purification mass spectrometry.

## SUPPORTING INFORMATION

Detailed methods to develop LAMP1-APEX iPSC lines; Additional figures for the fluorescence imaging of other APEX lines in neurons, optimization of protease amount, PCCA normalization, evaluation of false discoveries in KuB line, beads titration assay for other APEX lines, Venn diagram of all APEX proteomics and controls; Additional tables of interference peptide exclusion list, whole cell lysate protein list for H_2_O_2_ vs. PB vs. Ctrl groups, protein IDs from all APEX proteomics datasets, and known lysosomal protein coverage in three LAMP1-APEX probes.

## Notes

The authors declare no competing financial interest.

## ACKNOWLEDGEMENTS

The authors want to thank the Akos Vertes lab at GW for the access to the PEAKS software and Maia Parsadanian for technical assistance in cell culture. This project is financially supported by the GW Faculty Startup Fund. A.M.F acknowledges the Bourdon F. Scribner Fellowship from GW Chemistry Department. M.S.F. is supported by the National Institutes of Health (F30AG060722).

---

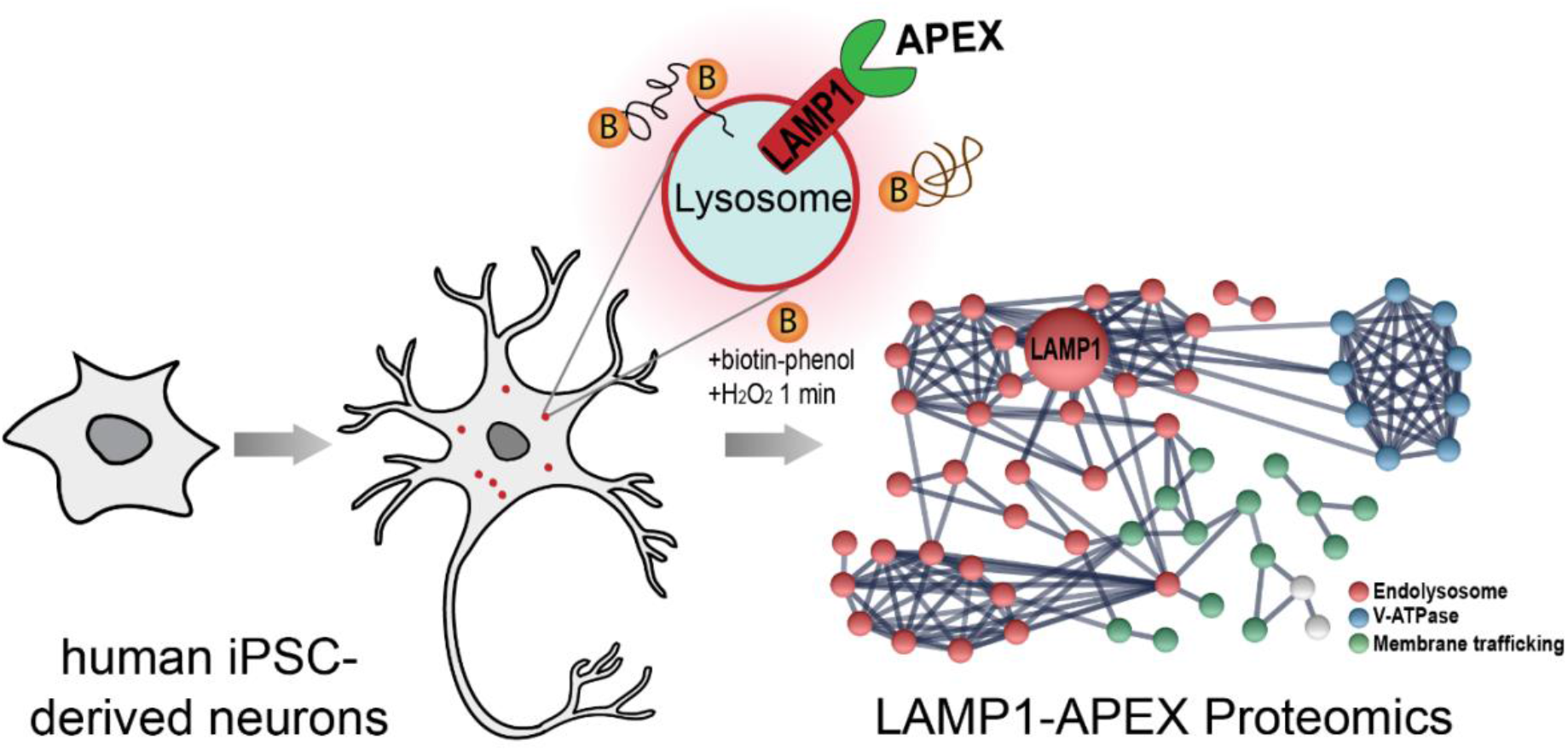
TOC Graphic.

