## Supplementary material for "Development and Comparative Evaluation of Endolysosomal Proximity Labeling-based Proteomic Methods in Human iPSC-derived Neurons": Supporting Information_Final.pdf

### TABLE OF CONTENTS


- Supplemental Methods: Development of LAMP1-APEX iPSC lines
- Figure S1. Fluorescence imaging of APEX activity in overexpression KuD-LAMP1-APEX and cytosolic NES-APEX iPSC-derived neurons.
- Figure S2. Distribution of peptide charges and precursor masses with different amount of proteases for on-beads protein digestion
- Figure S3: Scatter plots showing reproducibility between biological replicates in the same batch of APEX labeling experiment before and after normalization to PCCA.
- Figure S4. Evaluation of false discoveries in KuB-LAMP1-APEX using different control dataset.
- Figure S5. Beads titration assay for overexpressed APEX probes: KuB-LAMP1-APEX, KuD-LAMP1-APEX, and cytosolic NES-APEX.
- Figure S6. Venn diagram of all identified proteins from all APEX probes and controls.
- Table S1. Interference peptide exclusion list
- Table S2. Whole cell lysate protein list for $H_2O_2$ vs. PB vs. Ctrl groups (for Figure 6A)
- Table S3. Protein IDs from all APEX-proteomic datasets
- Table S4. Known lysosomal protein coverage in three LAMP1-APEX probes

#### Supplemental Methods: Development of LAMP1-APEX iPSC lines

For the endogenous KI-LAMP1-APEX line, iPSCs were engineered by CRISPR-mediated homologous recombination of the APEX2 transgene into the endogenous LAMP1 gene. APEX2 is the second generation of APEX with improved enzymatic activity.[1] We refer APEX2 as APEX in the paper for simplicity. Briefly, 1.5 million cells were seeded onto a 6-well dish for reverse transfection with Lipofectamine Stem (ThermoFisher). A ribonucleoprotein particle containing a crRNA targeting the 3' end of the LAMP1 ORF, tracrRNA, and recombinant Cas9 protein, was co-transfected with a custom DNA plasmid harboring 1-kb homology fragments flanking the APEX gene and a fluorescent selection cassette (Genewiz). The following day, the cells were dissociated onto a 10 cm dish, and maintained on Essential 8 medium for one week. Genomic DNA was collected from the unpurified cells using a Quick-DNA Microprep Kit (Zymo) and endogenous integration of APEX2 at the 3' end of a single LAMP1 ORF allele was confirmed by PCR. When the cultures reached an 80-90% confluency, a FACS Sony SH800S Cell Sorter was used to seed a 96-well plate with individual fluorescent cells, and scaled to 6-well dishes.

Two overexpression lines, KuD-LAMP1-APEX and KuB-LAMP1-APEX were generated to compare with the endogenous KI-LAMP1-APEX line. The HA line was generated by TALEN-mediated integration of a tetracycline-inducible KuD-LAMP1-APEX transgene at the CLYBL gene (UNIPROT: Q8N0X4). Since the high expression level of the TET-On promoter may drive partial mislocalization of LAMP1 to the cell membrane, we used a detuning strategy of upstream open reading frames (uORFs) to decrease transcriptional efficiency and enable more physiologic expression levels of LAMP1-APEX. Our previously developed LAMP1-APEX employed the moderate Kozak/uORF detuning strategy "KuB" (CAAATGGGTTGAACC-start).[2,3] Compared to the KuB line, KuD line employed the stringent Kozak/uORF "KuD" (GGGATGGGTTGATTT-start). KuB is predicted to reduce expression from the TET-ON promoter driving LAMP1-APEX to <15% of a consensus Kozak sequence (GCCACC-start), whereas KuD is predicted to reduce expression to <2% of the consensus sequence. The successful integration of APEX onto the LAMP1 locus was confirmed by PCR for each transgenic iPSC line.

### Supplemental Figures:

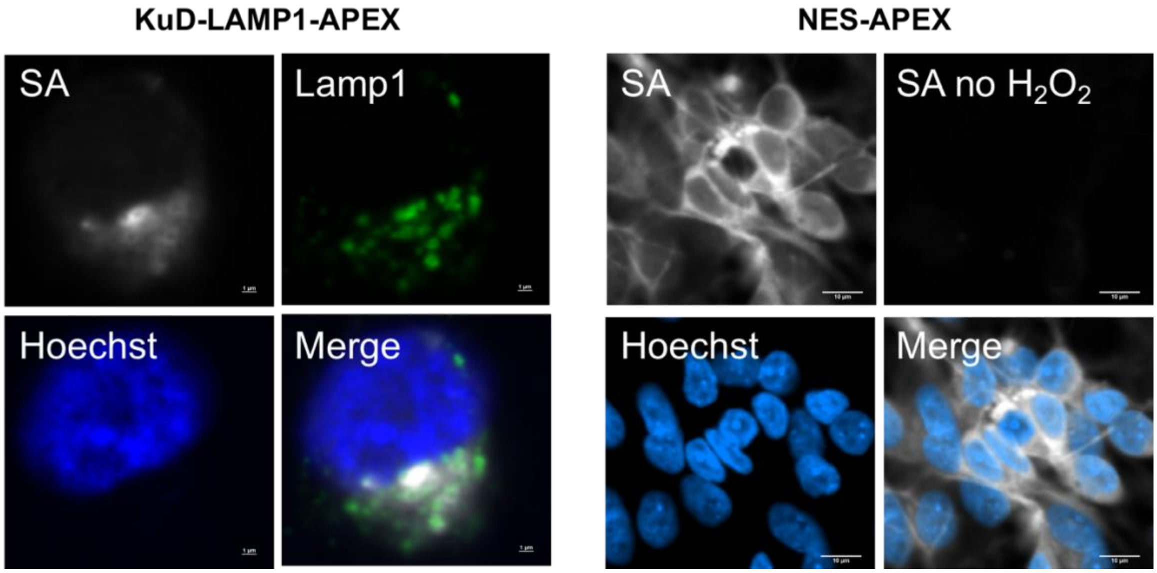


**Figure S1. Fluorescence imaging of APEX activity in overexpression KuD-LAMP1-APEX neurons (left) and cytosolic NES-APEX neurons (right).** Biotinylation is visualized by staining against streptavidin (SA) Fluor 680 (far red). Hoechst is a nuclear marker (blue). LAMP1 (green) is used as an endolysosome marker. Control neurons without $H_2O_2$ treatment exhibit no biotinylation signals. The APEX activity of the KuB-LAMP1-APEX probe was shown in our previous publication.[3]

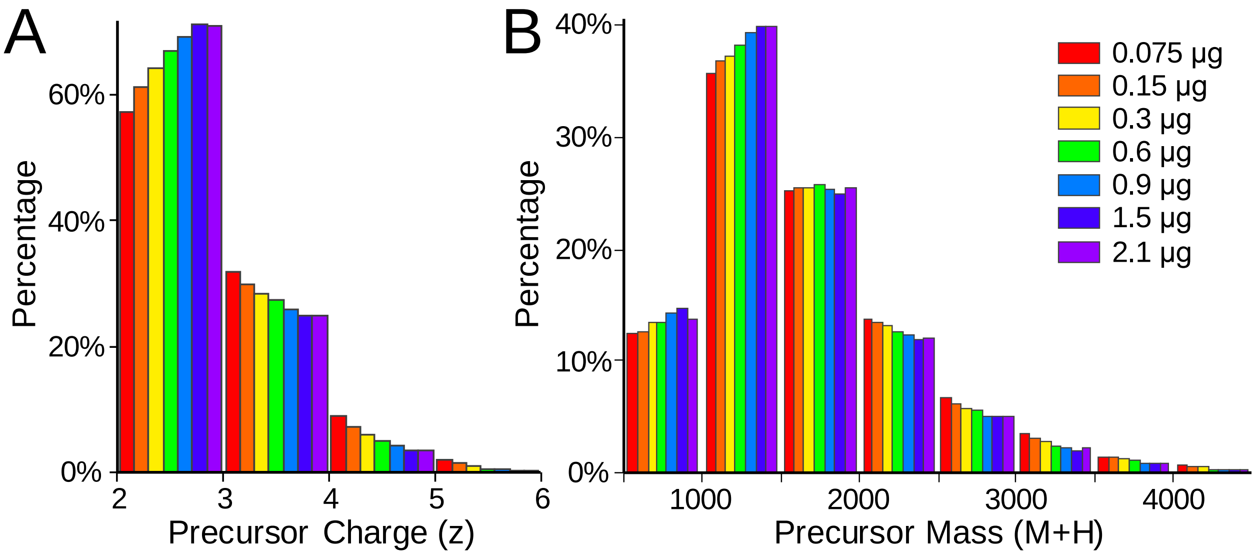


**Figure S2. Distribution of peptide charges (A) and precursor masses (B) with different amount of proteases for on-beads protein digestion.** Increased amount of protease (Trypsin/LysC mix) shifted the peptides towards lower charges and smaller precursor masses.

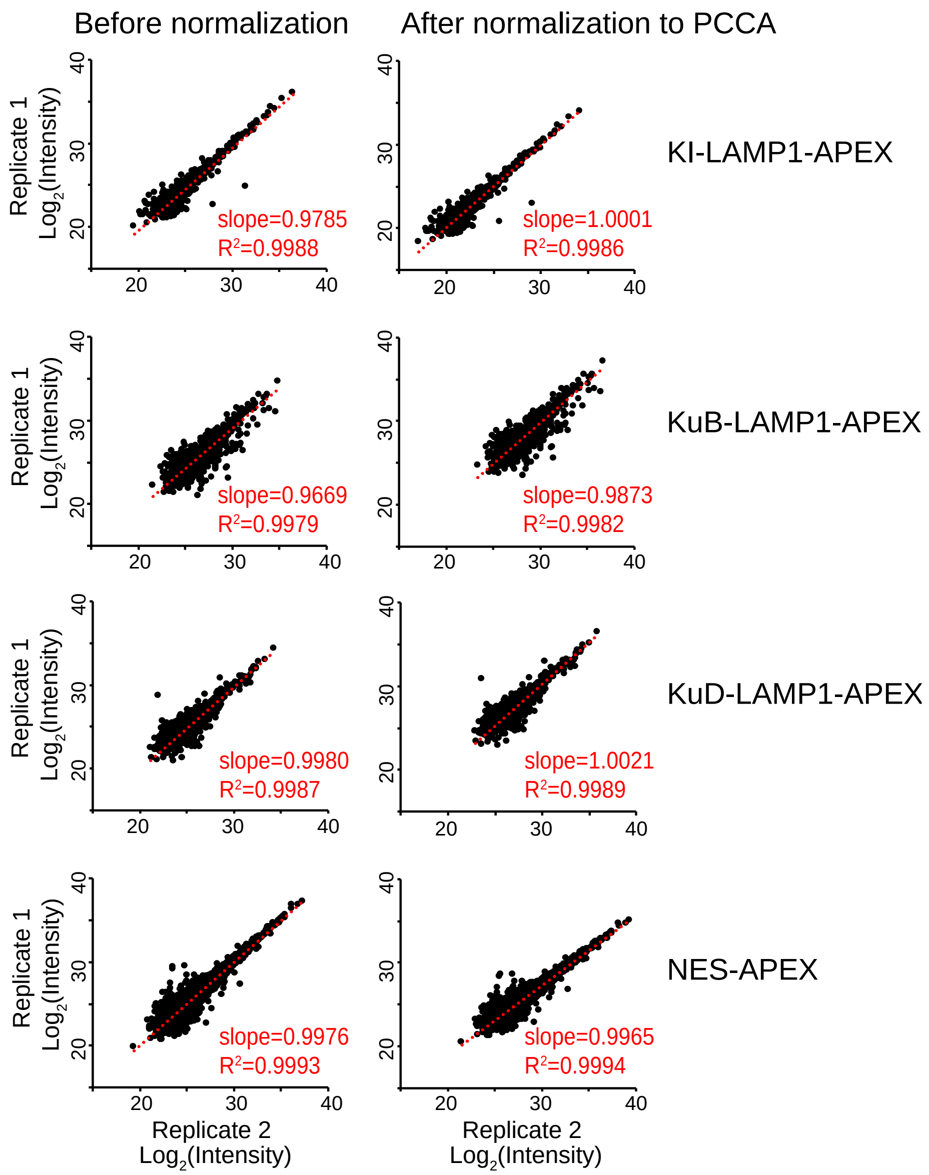


**Figure S3: Scatter plots showing reproducibility between biological replicates in the same batch of APEX labeling experiment before and after normalization to PCCA.**

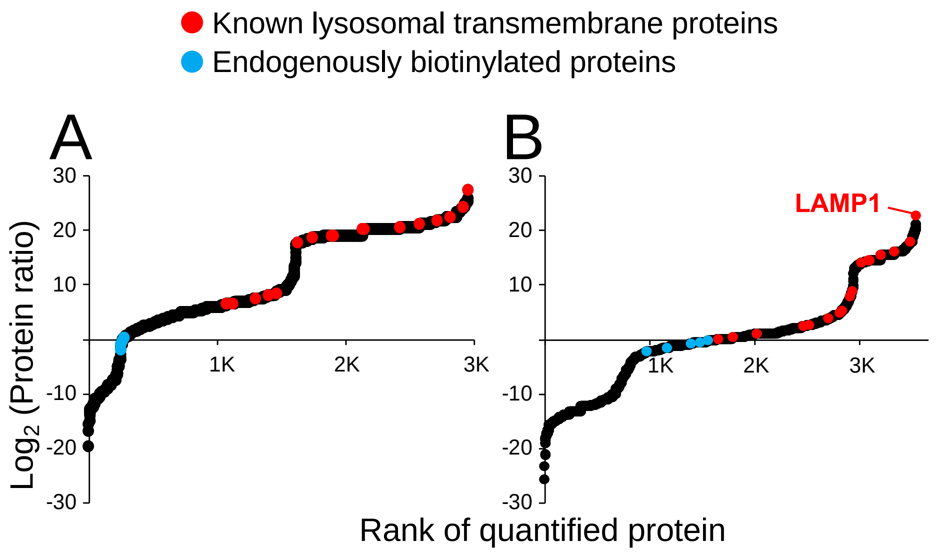


**Figure S4. Evaluation of false discoveries in KuB-LAMP1-APEX using different control dataset.** (A) KuB-LAMP1-APEX Proteomics with no-APEX line as control; (B) KuB-LAMP1-APEX with NES-APEX as control. Protein intensities were normalized to the most abundant endogenously biotinylated protein, PCCA.

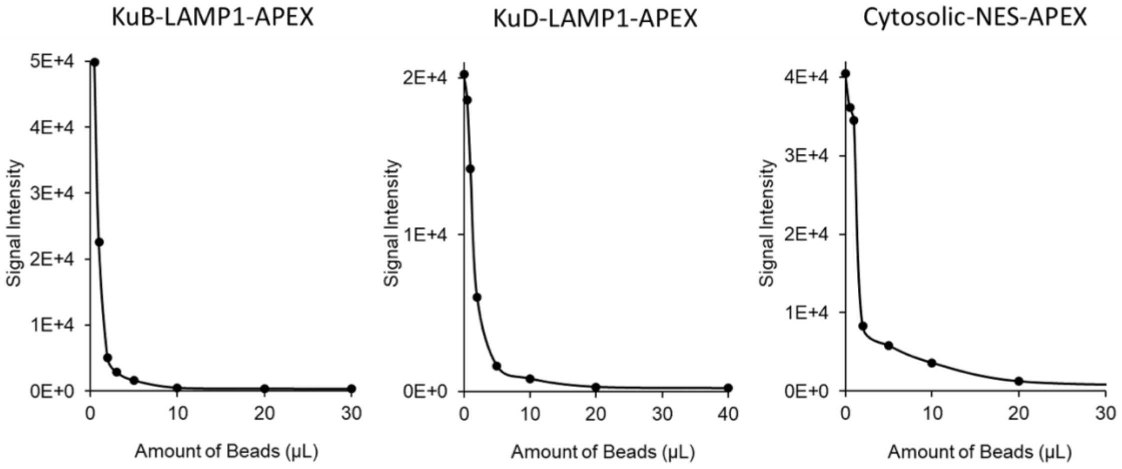


**Figure S5. Beads titration assay for overexpression APEX probes: KuB-LAMP1-APEX, KuD-LAMP1-APEX, and cytosolic NES-APEX.** Increasing amount of beads were incubated with 20 μg of input protein lysate in different tubes, followed by dot-blot assay against streptavidin staining.

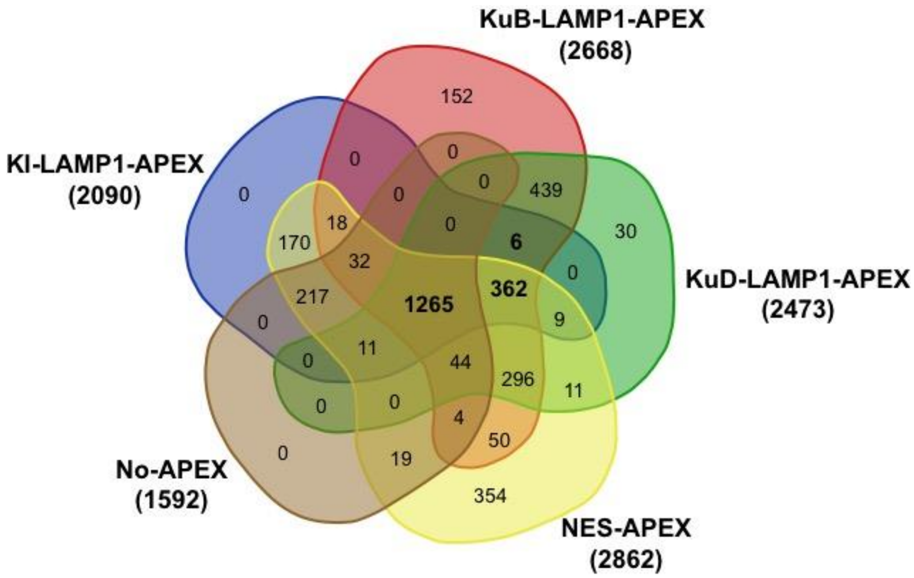


**Figure S6. Venn diagram of all identified proteins from all APEX probes and controls.** Three LAMP1-APEX proteomics, cytosolic NES-APEX, and No-APEX control groups.
